# Precision in a sea of profusion: Myelin replacement triggered by single-cell cortical demyelination

**DOI:** 10.1101/2019.12.16.877597

**Authors:** Nicolas Snaidero, Martina Schifferer, Aleksandra Mezydlo, Martin Kerschensteiner, Thomas Misgeld

## Abstract

Myelin—rather than being a static insulator of axons—is emerging as an active participant in circuit plasticity. This requires precise regulation of oligodendrocyte numbers and myelination patterns. Here, by devising a laser ablation approach of single oligodendrocytes, followed by *in vivo* imaging and correlated ultrastructural reconstruction, we show that in mouse cortex demyelination as subtle as loss of a single oligodendrocyte can trigger robust cell replacement and remyelination timed by myelin breakdown. This results in reliable reestablishment of the original myelin pattern along continuously myelinated axons, while in parallel profuse isolated internodes emerge on previously unmyelinated axons. Thus, in mammalian cortex, internodes along partially myelinated cortical axons are typically not re-established, suggesting that the cues that guide ‘patchy’ myelination are not preserved through cycles of de- and remyelination. In contrast, continuous ‘obligatory’ myelin shows remarkable homeostatic resilience with single axon precision.

## INTRODUCTION

Myelin is a multi-layered membrane sheath around vertebrate axons, which in the central nervous system (CNS) is generated by oligodendrocytes (OL). It increases conduction velocity and provides metabolic support for axons ^1–3^. While myelin was previously perceived as a stable and stereotypical structure ^4^, recent work shows that myelination patterns can be dynamic and highly variable—especially outside classical white matter tracts ^5, 6^. This suggests that myelin could support circuit plasticity and hence contribute to higher brain functions, such as learning and memory. For instance, changes in myelin content have been associated with task learning in the adult brain ^7–9^, and indeed such learning requires active myelination, as well as oligodendrocyte precursor cell (OPC) proliferation and differentiation ^10, 11^.

Central to this new paradigm of adaptive myelin plasticity is the unique myelination pattern described in mammalian gray matter, e.g. in the cortex. In contrast to the myelination patterns in white matter tracts, such gray matter myelin forms a complex sequence with myelinated and unmyelinated stretches resulting in a discontinuous pattern along both excitatory and inhibitory axons, albeit with preference to sheath the latter ^6, 12, 13^. Current data suggest that the editing of such ‘patchy’ myelination patterns involves the emergence of new OLs from non-myelinating precursors. For instance, the number of OLs increases until late adulthood in the cortex ^5, 14^, but OPC numbers remain stable, implying proliferation to match the generation of new myelinating OLs, at least in rodents ^15, 16^. Furthermore, BCAS1 positive OLs, which are transiently present during the active phase of myelination, have been identified in adult gray, but not white matter ^17^, reinforcing the idea of a ‘myelination reserve’ that could contribute to plasticity throughout adulthood. At the same time, there appears to be a steady, albeit subtle loss of OLs in the adult cortex. For instance, based on C^14^ incorporation, 2.5% of oligodendrocytes in human adult cortex are renewed per year ^18^. In mice, OL loss might amount to as much as 1.5% per month in the adult motor cortex ^19^. While such small losses might be unlikely to cause conduction blocks in white matter tracts, they could be detrimental to precise circuit computation, such as spike timing-based plasticity ^20, 21^.

These findings raise the question how cortical myelination patterns can be maintained *in vivo*, given the conflicting demands of participating in ongoing plasticity of neural circuits, while at the same time stemming progressive OL dropout and preserving prior patterns. Indeed, while the ‘patchy’ appearance of cortical myelin might hint towards a random pattern, the adaptation of myelin sheaths during circuit development and plasticity ^22^ suggests that regulatory principles exist to guide both initial cortical myelination, but also later spontaneous or induced additions of sheaths to cortical myelin. While histological and ultrastructural investigations of mammalian white matter tracts have supported the view that reformed myelin after demyelination can differ substantially from the original ensheathment, e.g. in internode length and thickness ^23, 24^, recent investigations of the developing zebrafish spinal cord show that primary myelination can resume with precise internode replacement if reset by OL ablation ^25^. However, as primary and secondary remyelination differ substantially ^26^, and highly regenerative zebrafish larvae lack a cortex or even a clear-cut white/gray matter divide ^27, 28^, it remains speculative, whether this new tenet of internode restoration applies to remyelination and homeostasis in mammalian cortex. Understanding this is particularly important in the context of demyelinating diseases like multiple sclerosis, where extensive cortical demyelination is emerging as a major pathological feature and a central target of remyelination therapies ^29–35^.

## RESULTS

### Laser ablation results in single oligodendrocyte loss with minimal off-target perturbation

Thus, to study the effect of losing a single OL on the complex myelin pattern in mouse gray matter, we utilized two-photon laser ablation ^25, 36^ in 3-6 month old *Plp*:GFP mice ^37^ followed by chronic intravital imaging through a cranial window above the somatosensory cortex ^38^. This ablation technique enabled us to monitor the physiological response to discrete myelin loss in an intact environment over 80 days (**Fig. 1a**). To exclude chronic off-target damage of our ablation paradigm, we used AAV-*Mbp*:mem-tdTom injections in *Cx_3_cr1^GFP/+^* knock-in mice ^39^, allowing simultaneous visualization of OLs and microglia. Laser ablation led to immediate loss of the OL cell body and a swift reaction of resident microglia ^40, 41^. Still, tissue damage was minimal, resulting in scar-free resolution of the microglial reaction within 24 hours (**Supplementary Fig. 1**). In addition, unperturbed axonal morphology was observed by correlative electron microscopy (EM) or sparse neuronal labeling after the loss of myelin in the OL ablation setting (AAV-*Mbp*:mem-tdTom injections in *Thy1*:GFP^M^ mice; ^42^, **Supplementary Fig. 2**). Thus, OL laser ablation allowed us to explore single cell repair strategies of myelin in a minimally altered CNS microenvironment.

**Fig. 1.**
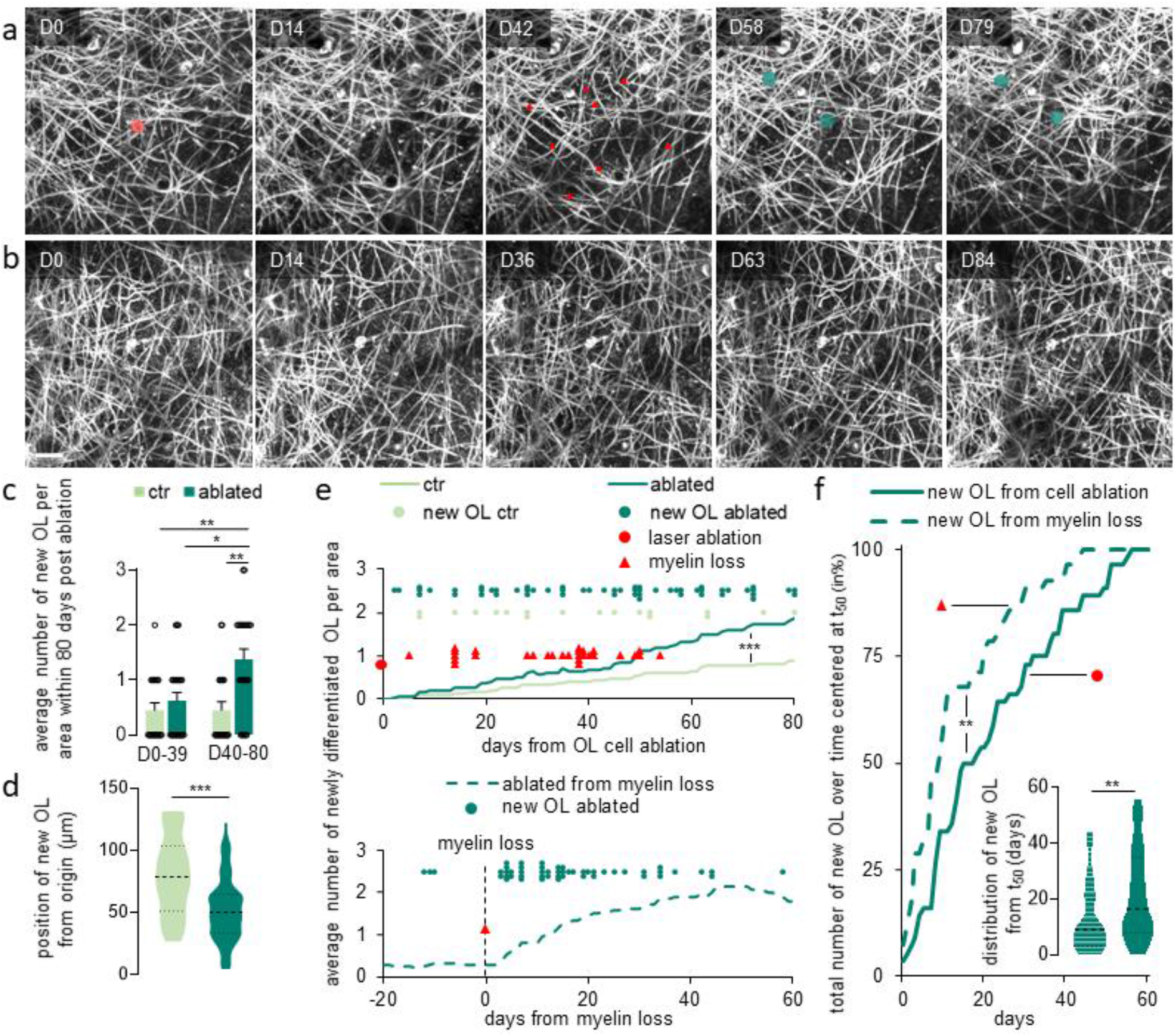
Demyelination due to single OL ablations triggers OPC differentiation in cortex. **a**: Longitudinal *in vivo* imaging of demyelination after single OL laser ablation (ablated cell highlighted in red at D0 and lost myelin segments marked by red arrowheads at D42) followed by new OL maturation and remyelination in *Plp*:GFP mice (new cells highlighted in green from D58). **b**: Longitudinal *in vivo* imaging of non-ablated control area. Display a-b: Maximum intensity projections of 30 μm. **c**: Average number of newly matured OLs per area (x/y/z: 203*203*60 μm^3^) over 80 days for control and ablated areas. Before D40 control: 0.5 ± 0.1; ablated: 0.7 ± 0.1; after D40 control: 0.5 ± 0.2; ablated: 1.4 ± 0.2 (control: n=20 and ablated: n=24 areas from n=10 and n=8 mice per group, respectively; mean ± SEM, Kruskal-Wallis with Dunn’s multiple comparison test). **d**: Average distance of the newly matured OL from the ablated cell or from the center of the control area a D0 (control: 78.9, n=17; ablated: 50.3μm, n=56; median, two-tailed t-test). **e, upper panel**: Timing and distribution of myelin loss and newly matured OL starting from ablation, imaged in ablated and in control areas, with curves representing the average cumulative number of new OL per area in ablated and control areas (Kolmogorov-Smirnov test). **e, lower panel**: Timing of newly matured OL centered on myelin loss in ablated areas, with curve representing the average cumulative number of new OL per area in ablated areas. **f**: Cumulative distribution of newly matured OL from the t_50_ values, obtained from the curves centered on the cell ablation or myelin loss (displayed in **e**) (Kolmogorov-Smirnov test). **Insert**: Variance of newly matured OL distribution timing from the t_50_ (from t_50_ of ablation: 16.5 days and from t_50_ of myelin loss: 9 days; n=56 new OLs from 24 ablated areas in 8 mice; median, Mann-Whitney U test). *** P<0.001, **P < 0.01, *P<0.05. Scale bar: 20μm.

### Timing of new oligodendrocyte appearance after ablation coincides with myelin loss

We performed long-term repetitive imaging following the ablation of mature cortical OLs (**Fig. 1**; **Supplementary Movie 1**). We observed loss of myelin and appearance of one to three newly matured OLs within 80 days after ablation in 95.8% of the studied areas (of these, one cell: 18.2%; two: 50.0%; three: 31.8%; **Fig. 1a** and **c**). In nearby non-ablated areas, the appearance of new OLs was less frequent (60%; one cell: 50%; two: 50%; three: 0%) at a baseline rate in line with previous reports ^5^. Accordingly, the average number of new cells was increased >3-fold by ablation (**Fig. 1b, c**; **Supplementary Fig. 3a**). This suggests that the removal of a single mature OL triggered the homeostatic differentiation of new myelinating cells. Indeed, we found these new OLs in spatial correlation to the ablation site within our field of observation: New OLs appeared significantly closer to the laser-targeting site in the center of field after ablation compared to ‘spontaneous’ new OLs in controls (**Fig. 1d**: **Supplementary Fig. 3b**).

We next monitored the timing of new OL emergence by repetitive intravital imaging. In the absence of laser ablation, new OLs were incorporated at a constant low rate in the adult cortex. However, starting 40 days after ablation, we observed a significant increase of newly matured OLs (**Fig. 1e**). The appearance of these induced new OLs did not seem to be tightly time-locked with the ablation itself, suggesting a different trigger. We noticed that ablation-induced myelin loss locally executed by microglial cells (**Supplementary Fig. 4**) was likewise variably delayed between 7 and 55 days post ablation (**Fig. 1e**), consistent with previous observations after genetic bulk removal of OLs ^43^. Thus, we asked, whether centering the emergence of new OLs on the myelin loss time-point, rather than the ablation, would yield a more uniform time course. Indeed, there was a clear increase in the rate of OL insertion that started within days after myelin loss (**Fig. 1e, f**). Notably the slope of the accumulation curve of new OLs was significantly steeper if it was time-locked on myelin loss rather than on OL ablation (**Fig. 1f**). This suggests that the appearance of the new OL was triggered by myelin degradation and exposure of denuded axonal surface, rather than OL death.

### Single oligodendrocyte ablation locally induces exuberant myelin formation

Next, we characterized the remyelination events following OL ablation. Our imaging technique allowed us to achieve single internode resolution *in vivo* for cortical volumes exceeding the typical territory of a single OL (**Supplementary Movie 1**) and to track internode fate after single OL ablation (**Fig. 2a**) or during spontaneous OL differentiation in control areas (**Fig. 2b**). First, we investigated the morphology and internodal territory of the new OL in relationship to the original ablated OL. While dense *Plp*:GFP labeling did not allow single OL reconstructions, subtraction of the pre- and post-myelin loss frames enabled reconstructing the ablated OL’s prior territory ^36^ and comparing it to the new cells’ morphology. Compared to the lost territory, the total regenerated myelin length produced by the newly matured OLs increased by 40%, as the total number of internodes was increased by about 60% compared to before (**Fig. 2c**). The number of internodes per OL was not different between the original and newly matured OLs, indicating that this increase in internode number after ablation is mainly due to the increased number of new OLs post ablation (**Fig. 1c**). Notably, while on average the internodes that were formed after cortical remyelination were shorter than the pre-existing ones (**Fig. 2d**), they were longer than the new internodes formed by spontaneously emerging OLs (**Fig. 2c-d**). This suggests that demyelination frees axonal surface that is especially supportive of myelination and induces longer internodes than normally formed by spontaneous myelination in adult cortex.

**Fig. 2.**
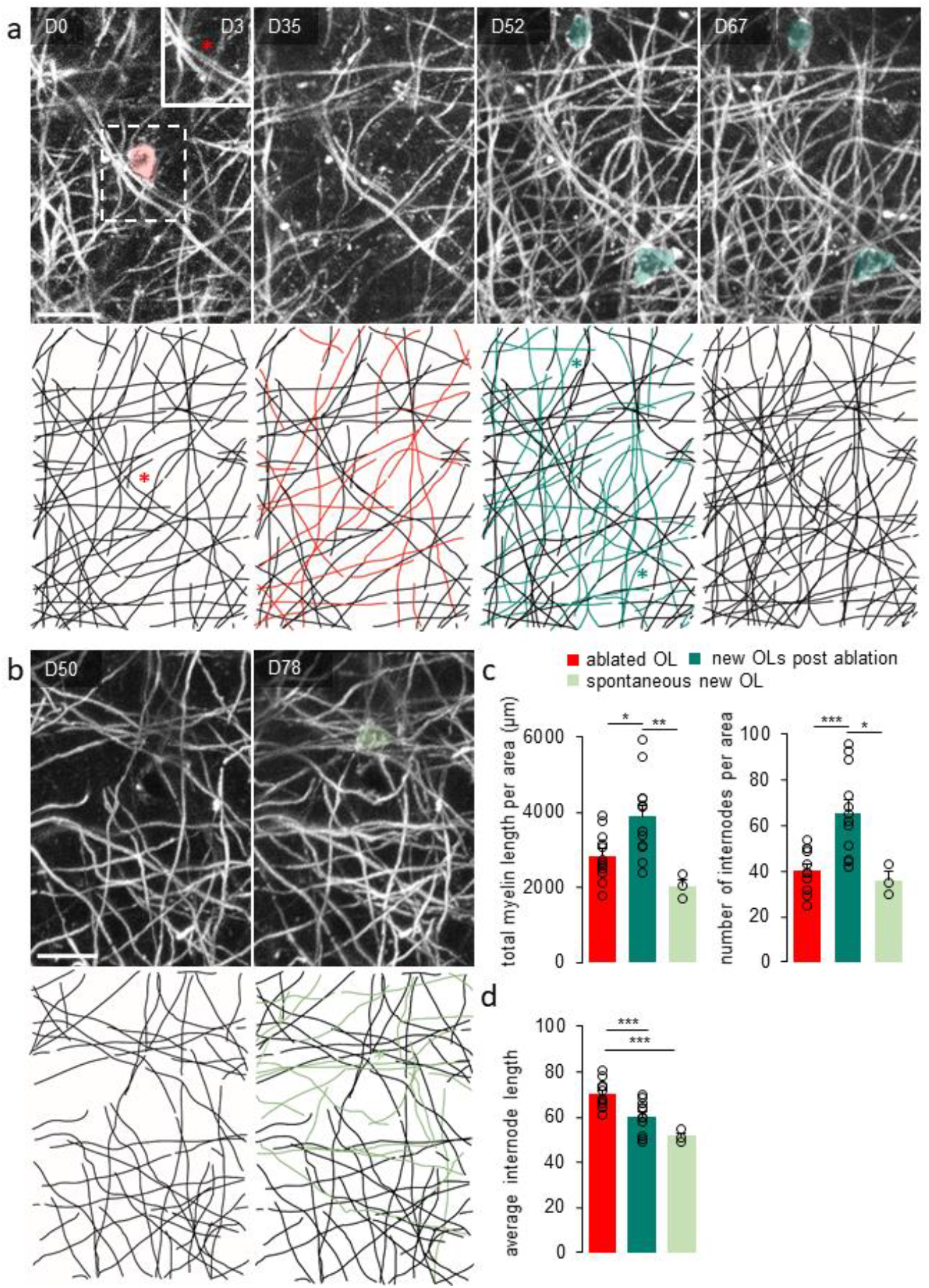
Single OL ablation induces excessive remyelination. **a**: Longitudinal *in vivo* imaging (upper panel) and manual tracing (lower panel) of an OL cell body and myelin loss (in red) at a single internode resolution after single OL ablation (at D0, red asterisk), followed by gained new OL differentiation and remyelination (in light green and marked by light green asterisks from D52). **Insert**: Imaging of the ablated site 3 days post OL ablation (red star indicates the position of the OL ablated at D0). **b**: OL cell body and myelin (in dark green and marked by dark green asterisk) gained after spontaneous OPC differentiation (D78 after the start of the experiment) in a control area. Display a-b: Maximum intensity projections of 27.5 μm. **c, left**: Total myelin length lost and gained after single OL ablation or gained after spontaneous OL differentiation per area (x/y/z: 203*203*60 μm^3^; ablated OL: 2831 ± 183 μm, new OL post ablation: 3872 ± 311 μm, spontaneous new OL: 2026 ± 181 μm). **c, right**: Average number of lost and gained myelin segments after single OL cell ablation or gained after spontaneous OL differentiation per area (ablated OL: 40.5 ± 3, new OL post ablation: 66 ± 5, spontaneous new OL: 36 ± 4). **d**: Average internode length of the lost and gained myelin segments after single OL cell ablation or gained after spontaneous OL differentiation per area (ablated OL: 70.4 ± 1.6 μm, new OL post ablation: 60 ± 2.0 μm, spontaneous new OL: 51.5 ± 1.6 μm). Statistics for c and d: per x/y/z: 203*203*60 μm^3^ area; ablated: n=12 and spontaneous: n=3 areas from 5 and 3 mice per group, respectively. Statistics: mean ± SEM, one-way ANOVA with Tukey’s multiple comparisons test. *** P<0.001, **P < 0.01, *P<0.05. Scale bars: 20μm.

To test this hypothesis directly, we needed to match internodes reliably before and after myelin loss and then match them to their ultrastructural correlate. Here, we built on a previous study, which showed that in mouse somatosensory cortex reconstructing less than 40 μm of axonal trajectory suffices (together with tissue landmarks) to discriminate a light microscopically identified axon from all its neighbors in correlated 3D EM volumes ^44^. As the average internode length exceeded this ‘ambiguity’ length limit (**Fig. 2d**), we could reliably follow single internodes degenerating after OL ablation and determine, whether a new internode had replaced a lost one (‘remyelinated’) or had formed on a previously non-myelinated axon stretch *(‘de novo’;* **Fig. 3a**). Using this approach, we found that about two thirds of original internodes (66.4 ± 2.9%) were restored. Comparing the morphological features of remyelinated and *de novo* internodes at the cellular and ultrastructural levels (**Fig. 3b-e**) revealed no apparent differences in myelin compaction and cytoplasmic spaces (**Supplementary Fig. 5**). Remyelinated internodes were substantially longer than the *de novo* segments, which were comparable in length to new segments produced by spontaneously emerging OLs (**Fig. 3b, c; Fig. 2d; Supplementary Fig. 6**). However, both *de novo* and remyelinated internodes had thinner myelin compared to preexisting internodes more than 40 days after remyelination, leading to an increased g-ratio (**Fig. 3e**). While previously myelinated axons thus induced longer internodes, this difference was not driven by axon caliber as internodes of both classes sheathed axons of similar diameter, which also did not differ from the axons sheathed by unaffected, stable myelin (**Fig. 3d**; **Supplementary Fig. 6**). Thus, myelination history rather than the particular geometry of targeted axons determines internode size.

**Fig. 3.**
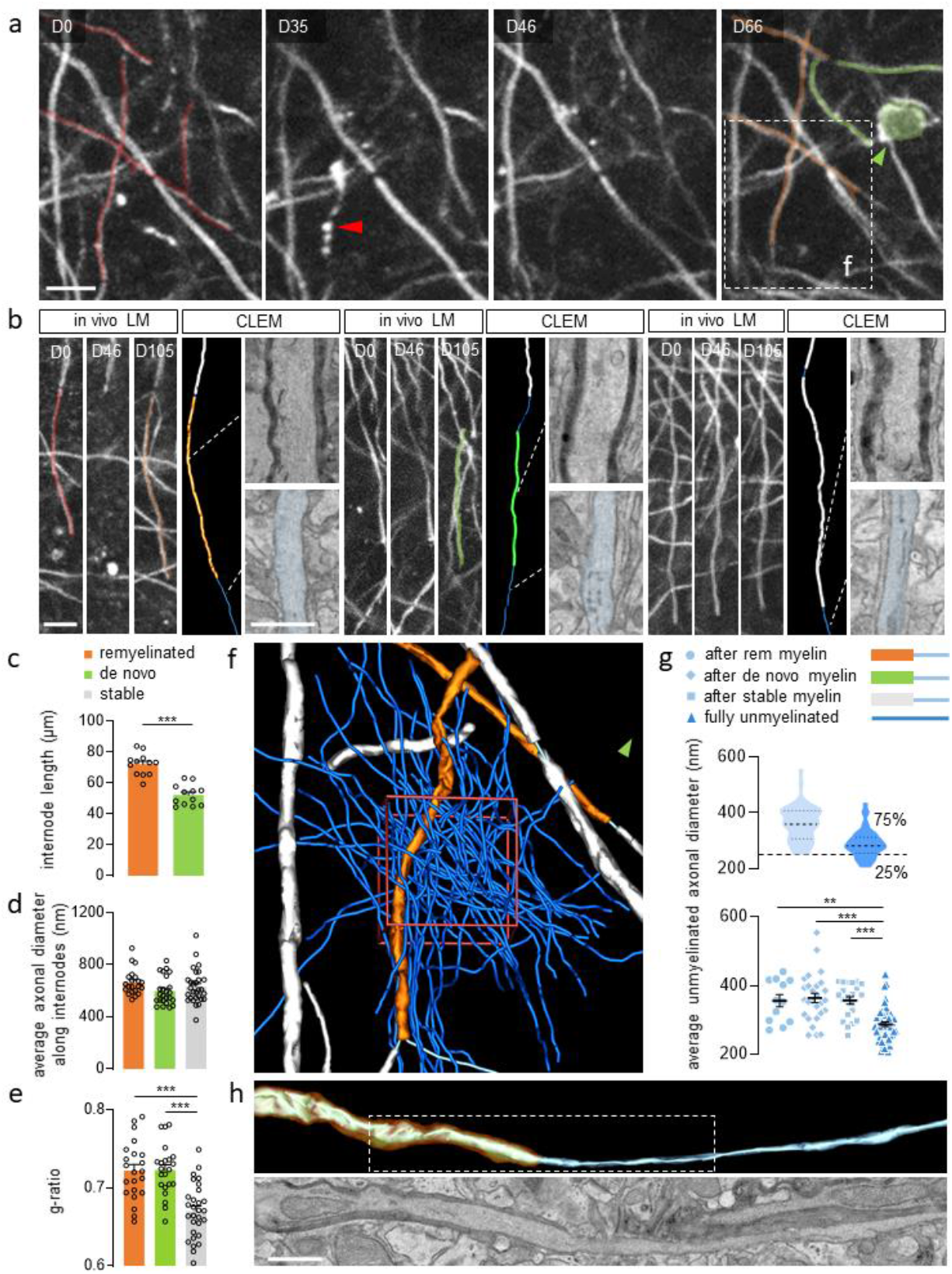
New OLs form internodes that closely resemble preexisting myelin. **a**: Representative images of longitudinal *in vivo* imaging allowing the identification of lost (red), degenerating (red arrowhead), remyelinated (orange) and *de novo* myelinated (green) segments after single OL ablation (newly matured OL: green arrowhead). **b**: Representative light microscopy images for each internode fate (remyelinated, *de novo* and stable) over the course of a single OL ablation experiment with correlated EM 3D reconstruction and high resolution scanning electron micrographs of the myelin sheath and underlying axon. Display a-b: Maximum intensity projections of 5 μm. **c**: Average length of remyelinated and *de novo* internodes (remyelinated: 72.1 ± 2.0 μm, *de novo:* 52.2 ± 1.9 μm; n=12 areas from 5 mice). **d:** Average axonal diameter of myelinated segments, measured along, for each fate: remyelinated: 664 ± 21,3 nm, *de novo:* 598 ± 21,9 nm or stable: 632 ± 24,3 nm (n=21 remyelinated, n=24 *de novo*, n=29 stable). **e**: Average g-ratio of remyelinated: 0.72 ± 0.008, *de novo:* 0.72 ± 0.007 and stable internodes: 0.67 ± 0.006 (n=21 remyelinated, n=24 *de novo*, n=29 stable). **f**: High-resolution EM reconstruction of unmyelinated axons ≥ 200 nm of diameter neighboring a remyelinated segment within 10*10*3 μm^3^ volume delimited in red (CLEM of area displayed in **a**). Green arrow indicates the location of newly matured OL. **g**: Distribution of unmyelinated axonal diameter, consecutive to a remyelinated segment: 356 ± 18,9 nm (n=12), *de novo*: 364 ± 13,7 nm (n=27) or stable: 357 ± 11 nm (n=21) internodes compared to fully unmyelinated axons within the boxed volume in f (288 ± 6,6 nm; n=52). **h:** Representative 3D EM reconstruction and corresponding electron micrograph illustrating axonal diameter transition at a remyelinated heminode. Statistics; mean ± SEM; **c**: two tailed t-test; **d,e**: one-way ANOVA with Tukey’s multiple comparisons test; **g:** Kruskal-Wallis with Dunn’s multiple comparisons test). *** P<0.001, **P < 0.01. Scale bars, light micrographs: 10 μm, electron micrographs: 1 μm.

### Remyelination in cortex is not random and is instructed by pre-ablation myelin patterns

The striking selectivity of the myelination pattern that was induced by OL ablation became apparent once we analyzed the immediate environment of the ablation-induced myelinating internodes in ultrastructurally resolved volumes, as this revealed the enormous density of potential myelination targets. We reconstructed 52 unmyelinated axons with a caliber ≥200 nm within 300μm^3^ (xyz: 10*10*3 μm^3^) surrounding a remyelinated segment (**Fig. 3f**). This represented about 10% of all axons in this volume ‘within reach’ of the new OL and suitable for myelination based on the previously described 200 nm threshold for CNS myelination ^44, 45^. We asked, whether these ‘ignored’ axons were simply systematically thinner than those chosen for induced myelination. To avoid the confounder that myelination *per se* can increase axon diameter (**Fig. 3h**) ^46, 47^, we compared the axon diameter distribution of fully unmyelinated axons with the distribution of the ‘naked’ axon parts that adjoined induced myelinated segments, as this likely represents the axon caliber at OL contact. Although the average diameter of the fully unmyelinated axons was indeed on average slightly smaller, we observed a 75% overlap between these two populations (**Fig. 3f, g**). We also found no difference in the naked axonal diameter consecutive to a remyelinated, *de novo* and stable internode, suggesting that also for these categories, axon diameter is not the driving force (**Fig. 3g**, **Supplementary Fig. 6**). Thus, the new OLs appear to be capable of a highly selective search for specific axons that are preferred targets for cortical remyelination.

To explore this specificity further, we related the myelination probability to the preexisting longitudinal myelination patterns along cortical axons. For this purpose, we classified internodes according to criteria that were phenomenologically defined in previous studies ^5^: ‘Continuous’, where an internode is adjoined by myelin on both sides, ‘interrupted’, with neighboring myelin on one side, and ‘isolated’ (**Fig. 4**). Based on three-dimensional reconstructions derived from our intravital imaging observations (**Fig. 4a, b**), we assigned the lost and gained internodes into these longitudinal axonal patterns (**Fig. 4c**). The degree to which the original myelin territory was restored was strongly influenced by the longitudinal pattern, into which these new internodes integrated: gaps in continuous myelination patterns torn by ablation were usually filled, while isolated segments were rarely replaced and interrupted patterns showed an intermediate repair efficiency (**Fig. 4d**). In accordance with this, when we broke down remyelinated vs. *de novo* new internodes, the former were predominantly part of continuous myelination patterns, while the latter were typically part of isolated patterns (**Fig. 4e**). Taken together these findings indicate that the apparently random pattern of cortical myelination contains sub-patterns that are resilient against demyelination and can be read out by emerging OLs with single internode precision. At the same time, these mechanisms do not operate along the majority of cortical axons with patchy myelination, which instead show profuse and exuberant de novo myelination, thus locally obscuring the original patterns.

**Fig. 4.**
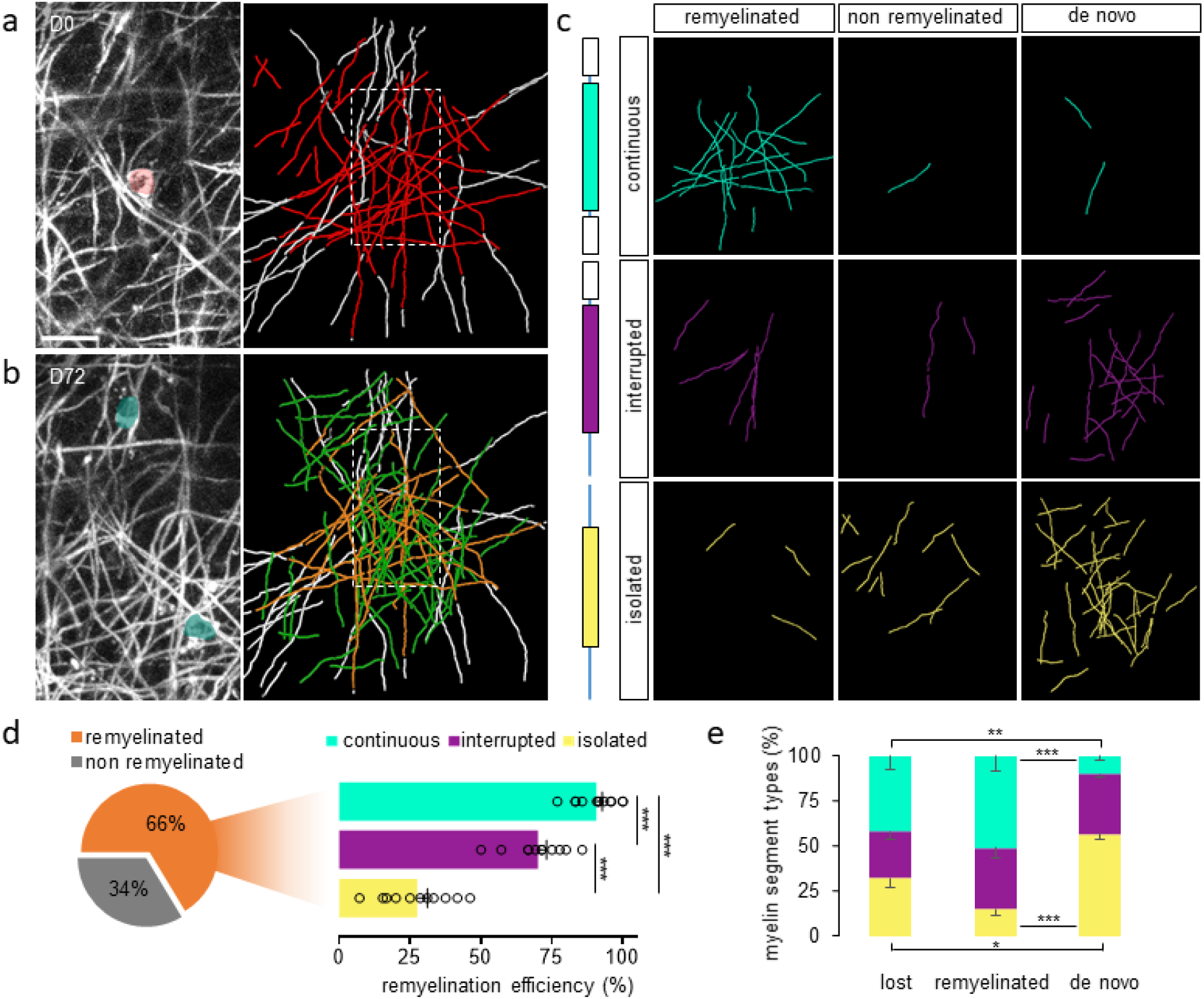
Cortical remyelination efficacy depends on the pre-ablation myelin patterns. **a, left**: Maximum projection of an OL prior to the laser ablation (in red at D0). **a, right**: 3D reconstruction of the lost myelin internodes between D0 and D38 (in red) and their neighboring internodes (in white). Dashed box represents the area displayed in the maximum projection. Red asterisk – position of the ablated OL. **b, left**: Maximum projection of a newly matured OL (in green at D72). **b, right**: 3D reconstruction of the gained myelin internodes between D38 and D72 (remyelinated segments in orange and *de novo* segments in green) and the neighboring internodes of the lost segments between D0 and D38 (in white). Dashed box represents the area displayed in the maximum projection. Green asterisks – position of the new OL. Display a-b: Maximum intensity projections of 30 μm. **c**: Traces of internodes separately displayed according to their post-ablation origin (remyelinated, non-remyelinated and *de novo*) and their pattern types (continuous, interrupted and isolated) after single OL ablation. **d**: Total and type specific remyelination efficiency from the single OL laser ablation (total: 66.4 ± 2.9%; continuous: 90.7 ± 2.1%, interrupted: 70.1 ± 3.1% and isolated: 27.5 ± 3.6%; n=12 areas from 5 mice, mean ± SEM, one way ANOVA with Tukey’s multiple comparisons test). **e**: Internode type distribution of the lost, remyelinated and *de novo* myelin fractions (n=10 areas from 5 mice, mean ± SEM, Kruskal-Wallis with Dunn’s multiple comparisons test). *** P<0.001, **P < 0.01. Scale bar: 20 μm.

## DISCUSSION

The adult cortex is characterized by a pattern of myelination that at first glance appears random, but that has been suggested to be specific and functionally important ^6, 12, 13, 48^. We now demonstrate that even loss of a single OL from this pattern can trigger a remarkably local and stereotypical response—during which one to three newly matured OLs replaced the original cell. These new OLs then remyelinate two thirds of the original myelin sheets, with a strong bias towards axons that originally were fully myelinated (where >90% of lost sheets are replaced), as opposed to the isolated internodes that were rarely remyelinated (<30%; **Fig. 4**). At the same time, many axons in the surrounding receive profuse *de novo* myelination by isolated internodes, which overcompensate the lost isolated internodes of the original cell and result in a local 40% excess of discontinuous myelination. Thus, the cortical myelin pattern harbors two sub-patterns of drastically differing resilience against small demyelinating perturbations: One—along continuously myelinated axons—that is reestablished with remarkable single internode precision vs. another consisting of isolated internodes that is not reestablished, but replaced by profuse and excessive *de novo* myelination frequently of previously unmyelinated axons. This duality allows remyelination—on the one hand—to preserve some functionally important structural engrams, and—on the other hand—to provide new imprints of ongoing plasticity or a substrate for later remodelling. Thus, cortical myelin is imbued with resilience, as well as plasticity, when confronted with physiological or pathological myelin turnover.

While it has long been known that demyelination is one of the few features of CNS injury that can be spontaneously repaired even in humans, the mechanisms of remyelination have remained controversial ^26^, and how precisely such replacement would reestablish the original pattern of myelination was unclear. This question has gained urgency, because increasing evidence suggests that myelin, especially ‘patchy’ cortical myelin, supports neuronal network plasticity during learning ^7–11, 49, 50^ using changes in number, thickness and spacing of internodes to regulate spike timing ^22, 51^. If myelination patterns indeed contain imprints of past plasticity, major demyelination insults—as they occur in demyelinating conditions such as multiple sclerosis that prominently affect the cortex ^52^ —might thus erase such myelin engrams at least transiently. Can these prior patterns be reestablished by endogenous or therapeutically supported remyelination, and if so, how precisely? Answering this question is not easy in injury paradigms where most OLs are damaged, as axon loss, glial scaring and smoldering inflammation confound following remyelination ^43, 53, 54^.

We therefore established a cortical laser ablation approach that can delete individual OLs with minimal perturbation of the CNS environment. Previously such ablations were used in mice to explore peripheral myelinating cells ^36^ and in developing zebrafish to study primary spinal cord myelination ^25^. In zebrafish spinal cord, precise reestablishment of internodes was observed, irrespective of the surrounding myelination pattern. However, these results do not seem to apply to mammalian cortex. While our results corroborate that a subset (90,7%) of cortical axons maintains the cues needed to reestablish their (continuous) myelination, the majority (71.2%) of discontinuously myelinated axons do not. Moreover, even those that remyelinate their original axonal targets do not typically re-establish a precise positional pattern of the internodes (**Supplementary Fig. 7**), resulting in only 8.2% of isolated internodes and 14.5% of internodes that integrate into a continuous pattern, reestablish internode position, as well as presence. Together with the altered g-ratio (**Fig. 3**), This indicates that in mammalian gray matter, axonal imprints of previous partial myelination are not reliably preserved through a cycle of oligodendrocyte loss and replacement.

Overall, our experiments reveal that while OL numbers and continuous myelination patterns are resilient, the detailed geometry of internodes is not. This implies that ‘classic’ functions of myelin, such as saltatory conduction and metabolic support are likely restored, but timing of action potentials might not. Moreover, three principles govern the replacement of lost OLs and the parallel emergence of precise and profuse remyelination: First, the trigger of OL differentiation is not the death of the ablated OL itself (which was cleared within 24 hours of ablation), but rather the following demyelination, which was delayed by as many as 50 days (average ~32 days). This is in line with other OL ablation paradigms, where myelin—which also on the protein level has an extremely long half-life ^55^—can survive long after OLs have died ^43^. In our experiments, we observed that the OL processes resealed rapidly after cell body ablation (**Supplementary Fig. 1**), then maintained their morphology for weeks before removal by resident microglia (**Supplementary Fig. 1 and 4**), which was consistently followed by OL replacement typically within 2 weeks (t_50_=9 days). Thus, it appears that myelin removal, which exposes the axonal surface, triggers OL maturation and remyelination—supporting the notion that myelin removal is a controlled process that can initiate subsequent repair ^56, 57^.

Second, while there was substantial overproduction of OLs and myelin, each individual new OL finally established a morphology—e.g. number of internodes—that on average matches that of the preexisting OL population. This suggests that in addition to extrinsic (e.g. axonal) cues that guide OL maturation and resilience of myelination patterns, there is also an OL-intrinsic component. At the same time, ablation resulted in a local excess of ‘isolated’ internodes compared to the original state, as in many cases there was overproduction of OLs. The significance of these exuberant internodes could be twofold. On the one hand, they could simply represent a random excess due to the OL’s ‘drive’ to myelinate, as also evidenced by the propensity of OLs to myelinate inert substrates in the absence of alternatives ^58^. This would imply that the myelinated axons are not in need or even especially suitable for myelination, but are ‘off-target’. Indeed, these isolated internodes are shorter than remyelinated ones, perhaps due to the lack of positive signals from the underlying axons or an increased density of topographical ‘barriers’ that prevent internode elongation, such as synapses and branch points ^6, 12, 13^. On the other hand, the excess internodes could be the imprint of the OL’s search for the ‘right’ axons that require myelination. This exuberance could still serve as the substrate for later plasticity-related pruning, as internode retractions appear to be a common occurrence in plastic myelin ^59^. However, on the time-sale of our follow-up (on average 40 days), pruning of fully differentiated internodes was not apparent. In either case, the stepwise increase in local myelin amount after OL loss could contribute to the increasing density of cortical myelin across adult life. Both in humans and rodents, OL turnover is a physiological occurrence ^18, 19^—and while the question of whether cortical myelination ever gets continuous on all myelination-receptive axons is still open, there is little doubt that myelination indeed progresses in the adult cortex, reaching up to 80% coverage on some axons ^14^.

Third, our data show that maturing OLs in the mature cortex have an uncanny ability to find efficient myelination targets in a vast ‘ocean’ of axons. Indeed, recent connectomics reconstructions of cortex showed that 220 axons traverse within a volume of 125μm^3^ ^44^ when the volume covered by a typical layer 1 OL alone is substantially larger (~ 5000-fold) than this^19^. Our own ultrastructural volume reconstructions corroborate this view, showing that a remyelinating OL ignores an enormous number of potential axonal targets, even if their diameters are intrinsically suitable for myelination. This is remarkable in light of the above cited fact that OLs readily myelinate inert fibers of suitable diameter *in vitro* ^58, 60^. It also implies that there likely are three populations of cortical axons that new OLs manage to differentiate: A few axons that require obligatory myelination (and as a result reestablish their continuous myelination pattern), many axons that actively suppress myelination and remain unmyelinated ^61^, and the remaining group of discontinuously myelinated axons that do not strongly enforce either outcome and as a result commonly change their myelination pattern after a cycle of de- and remyelination. The later finding implies that the dominant ‘patchy’ pattern of cortical myelination in mammalian cortex cannot be preserved even after minimal OL loss - suggesting that little long-term information storage can happen in these patterns and that such information would be comprehensively lost after pathological demyelination, even if remyelination ensues.

Overall, out data suggest that in the apparently random pattern of cortical myelin both highly regulated, but also spurious sub-patterns exist, which differ profoundly in their resilience even against minimal demyelination. Thus, our results reveal the degree of fidelity with which remyelination can naturally happen in mammalian cortex. This is critical information for developing efficient, but at the same time sufficiently precise therapeutic interventions to reverse pathological cortical demyelination ^30^.

## ACKNOWLEDGMENTS

We thank M. and N. Budak, as well as S. Taskin for animal husbandry; Y. Hufnagel, K. Wullimann and M. Schetterer for technical and administrative support. We thank B. Zalc (ICM Paris) for providing *Plp*:GFP mice, M. Terasaki (U. Connecticut), J. Lichtman and R. Schalek (Harvard U.) for EM advice and materials, and acknowledge Drs. L. Godinho, O. Karpiuk, T. Czopka and M. Simons for reading an earlier version of this manuscript. N.S. is supported by the Deutsche Forschungsgemeinschaft (DFG; Sn 149/1-1). Work in M.K.’s laboratory is financed through grants from the DFG (TRR128, Project B10 and B13), the European Research Council (ERC) under the European Union’s Seventh Framework Program (FP/2007-2013; ERC Grant Agreement n. 310932), the German Federal Ministry of Research and Education (BMBF; Competence Network Multiple Sclerosis), the “Verein Therapieforschung für MS-Kranke e.V.” and the German National MS Society (DMSG). Work in T.M.’s lab is supported by the DFG (CIPSM EXC114, CRC870, Mi 694/8-1, FOR 2879) and the ERC (FP/2007-2013; ERC Grant Agreement n. 616791). M.K. and T.M. are further supported by the DFG through a common grant (Ke 774/5-1/Mi 694/7-1), the Gemeinnützige Hertie Stiftung and the Munich Center for Systems Neurology *(SyNergy* EXC 2145), which also supports M.S.. N.S., M.S. and T.M. are additionally supported by the German Center for Neurodegenerative Diseases (DZNE).

## AUTHOR CONTRIBUTIONS

N.S., M.K. and T.M. devised the study. N.S. developed imaging technology, performed and analyzed two-photon imaging experiments. A.M. designed and cloned viral vectors. N.S. and M.S. established electron microcopy technologies, as well as generated ultrastructural data. N.S. analyzed the EM data. N.S., M.K. and T.M. wrote the paper with input from all authors.

## SUPPLEMENTARY FIGURES AND MOVIES

**Supplementary Figure 1.**
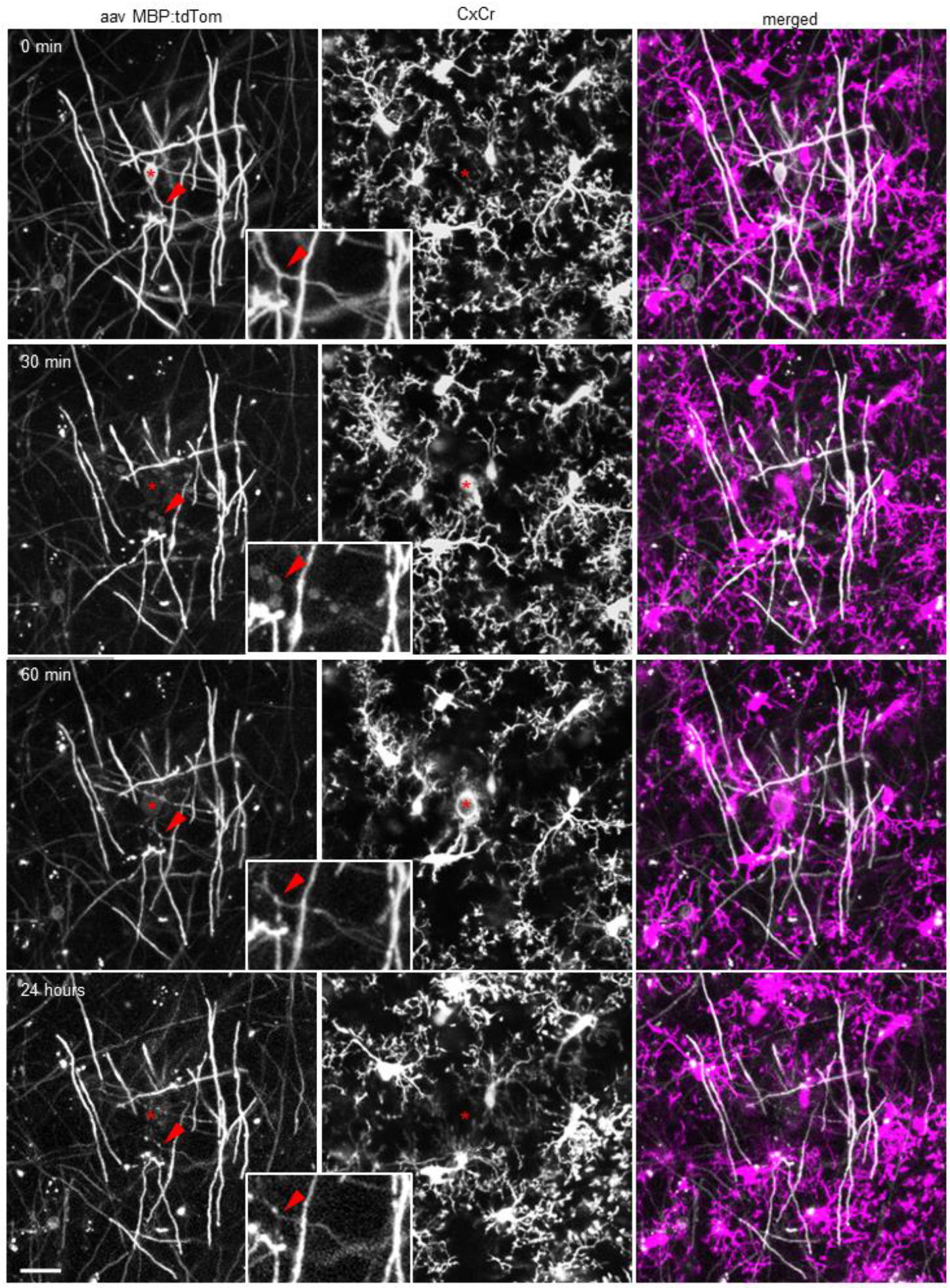
Microglial response to a single OL laser ablation. Intravital imaging of a cortical OL laser ablation with labelled resident microglial cells (*Cx_3_cr1*^GFP/+^ mouse). Cortical OLs are labelled by injection of AAV-*Mbp*:mem-tdTom. 0 minutes illustrate the pre ablation state. Thirty minutes post ablation, neighboring microglial cells start polarizing toward the ablated OL and by 60 minutes the cell body is surrounded by the neighboring microglial end feet (red star designate the ablated OL soma position). 24 hours post ablation no microglial scar was observed, and the ablated cells internodes persisted with preserved connecting processes (red arrowheads, **inserts**). Display: 10 μm maximum intensity projection. Scale bar: 20μm.

**Supplementary Figure 2.**
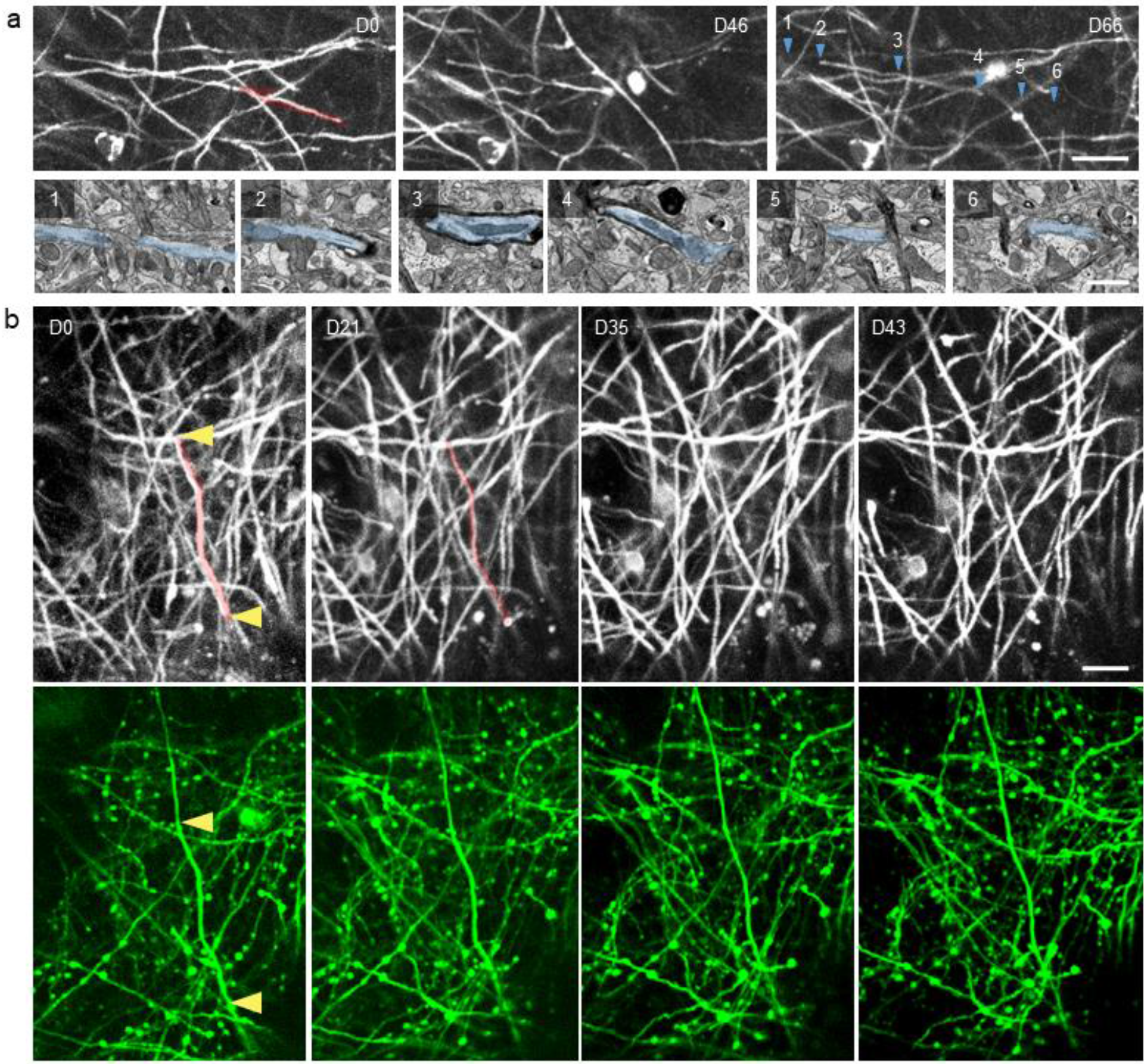
Unaltered axonal morphology after single OL ablation and demyelination. **a, upper panel**: Longitudinal *in vivo* imaging of a demyelination event without remyelination (highlighted in red). **a, lower panel:** Correlative electron micrographs of the partially myelinated axon (blue) displayed in the upper panel. The unmyelinated (1), nodal (2, 4), internodal (3) and non remyelinated (5, 6) portions of the axon are displayed from the areas pointed out at D66 (blue arrowheads). Display: 2 μm maximum intensity projection. **b**: Longitudinal *in vivo* imaging of axons near the OL ablation site until demyelination (**upper panel**: myelin labeling by injection of AAV-*Mbp*:mem-tdTom; **lower panel**: sparse axonal labeling in the *Thy1*:GFP^M^ line). The lost myelin segment is highlighted in red and the nodal regions by yellow arrowheads. Display: 2 μm maximum intensity projection. Scale bar, light micrographs: 20 μm, electron micrographs: 1 μm.

**Supplementary Figure 3.**
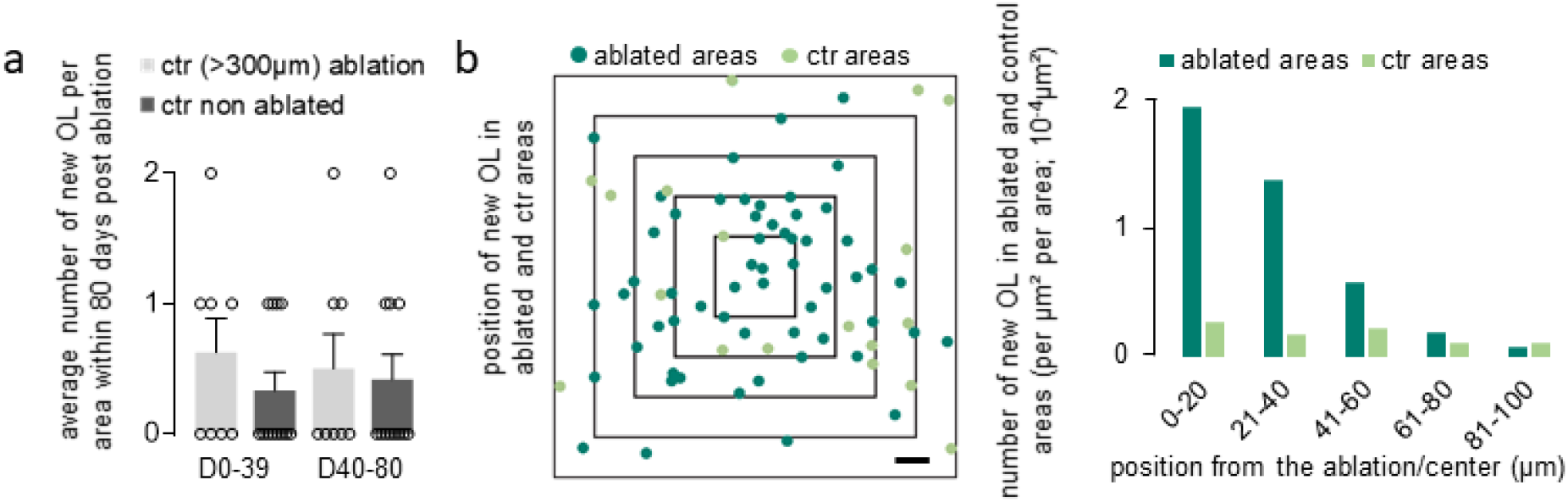
Spatiotemporal and morphological features of newly matured OLs. **a**: Average number of newly matured OL per area over 80 days for control areas in animals where laser ablation was performed—ctr ablated— (areas centered >300 μm away from the ablated cells) and in control animals where no OL ablation was performed—ctr non ablated—(ctr ablated: n=8 and ctr non ablated: n = 12 areas from 5 and 2 mice per group, respectively; mean ± SEM, Kruskal-Wallis with Dunn’s multiple comparisons test). **b, left**: Position of newly matured OLs in ablated and control areas centered on the ablated cell or the center of the field of view, respectively, at D0. **b, right**: Distribution of newly matured OLs per μm^2^ per area. Scale bar: 20 μm.

**Supplementary Figure 4.**
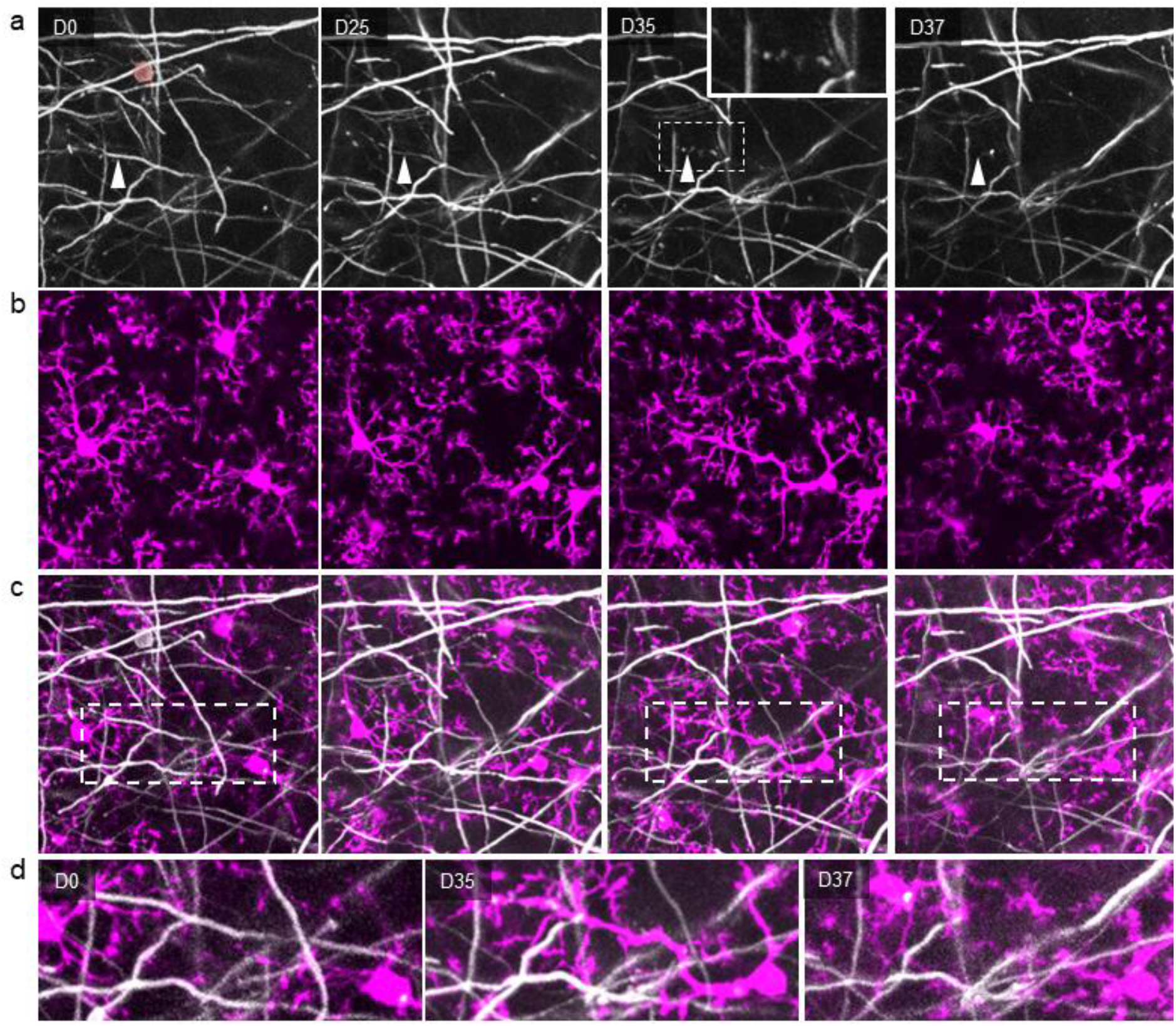
Secondary removal of myelin fragments by neighboring microglia. **a**: Longitudinal *in vivo* imaging of demyelination after single OL laser ablation (in red), highlighting a myelin sheath (white arrowhead) that is stable up to D25 post ablation, but shows degradation profiles at D35 (insert) and no labelling at D37 (myelin labeling: AAV-*Mbp*:mem-tdTom). **b**: resident microglia surrounding the lost myelin sheath (*Cx_3_cr1*^GFP/+^ mouse), **c**: merged channels. **d**: high magnification of the inserts in c, showing the transient alignment of microglial process with degenerating myelin at D35. Display: 10μm maximum intensity projection. Scale bar: 20μm.

**Supplementary Figure 5.**
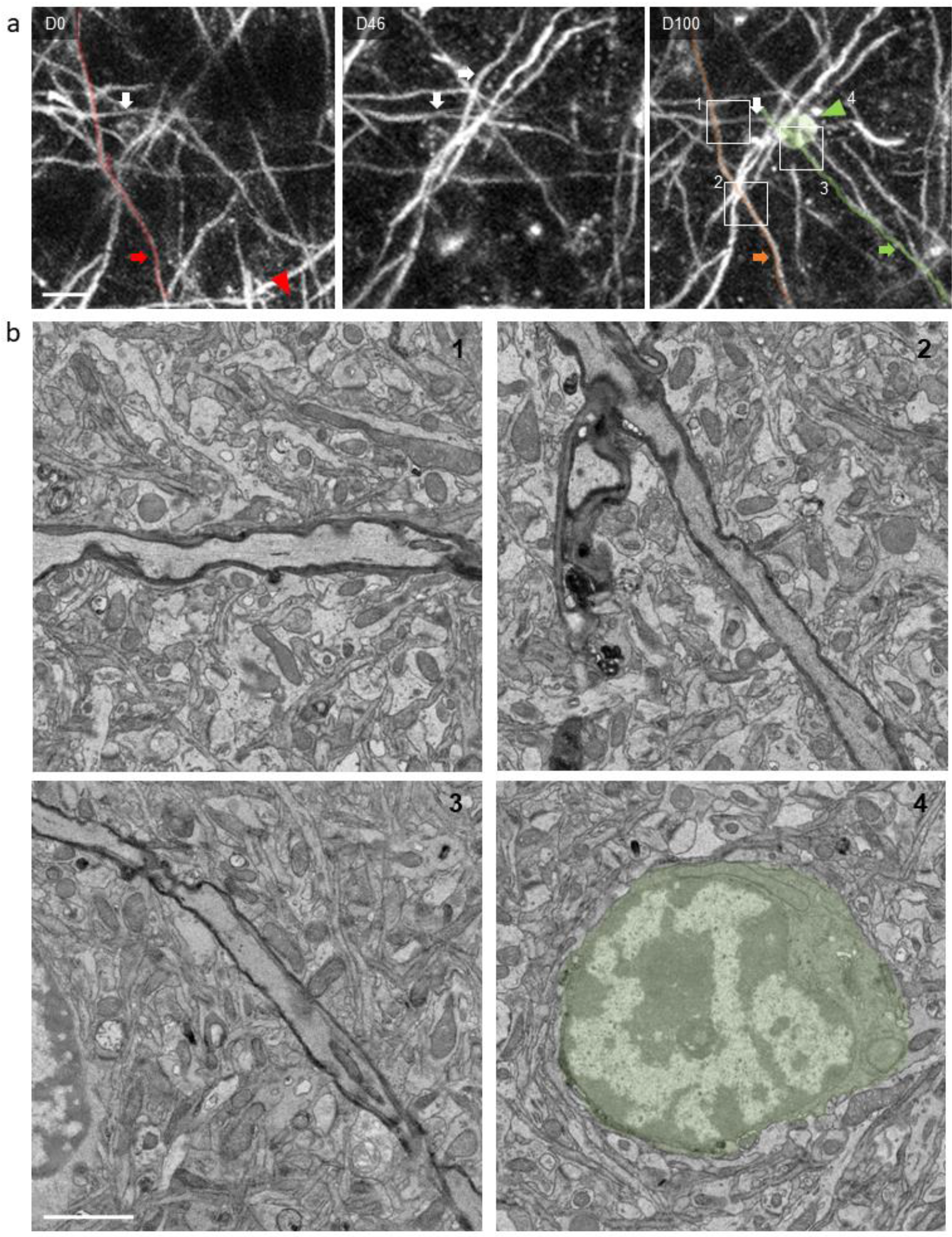
Correlative long-term intravital imaging with high-resolution volume electron microscopy. **a**: Longitudinal intravital imaging allowing the identification of the lost (red), remyelinated (orange), *de novo* myelinated (green) and stable (white) segments after single OL ablation (green arrowhead points towards newly matured OL). **b:** Correlative EM at D100 of the three areas marked in a, illustrating the ultrastructure of the different internode types (1: stable, 2: remyelinated and 3: *de novo*) and the newly matured OL soma (4: green). Scale bars: 10μm in **a** and 2 μm in **b**.

**Supplementary Figure 6.**
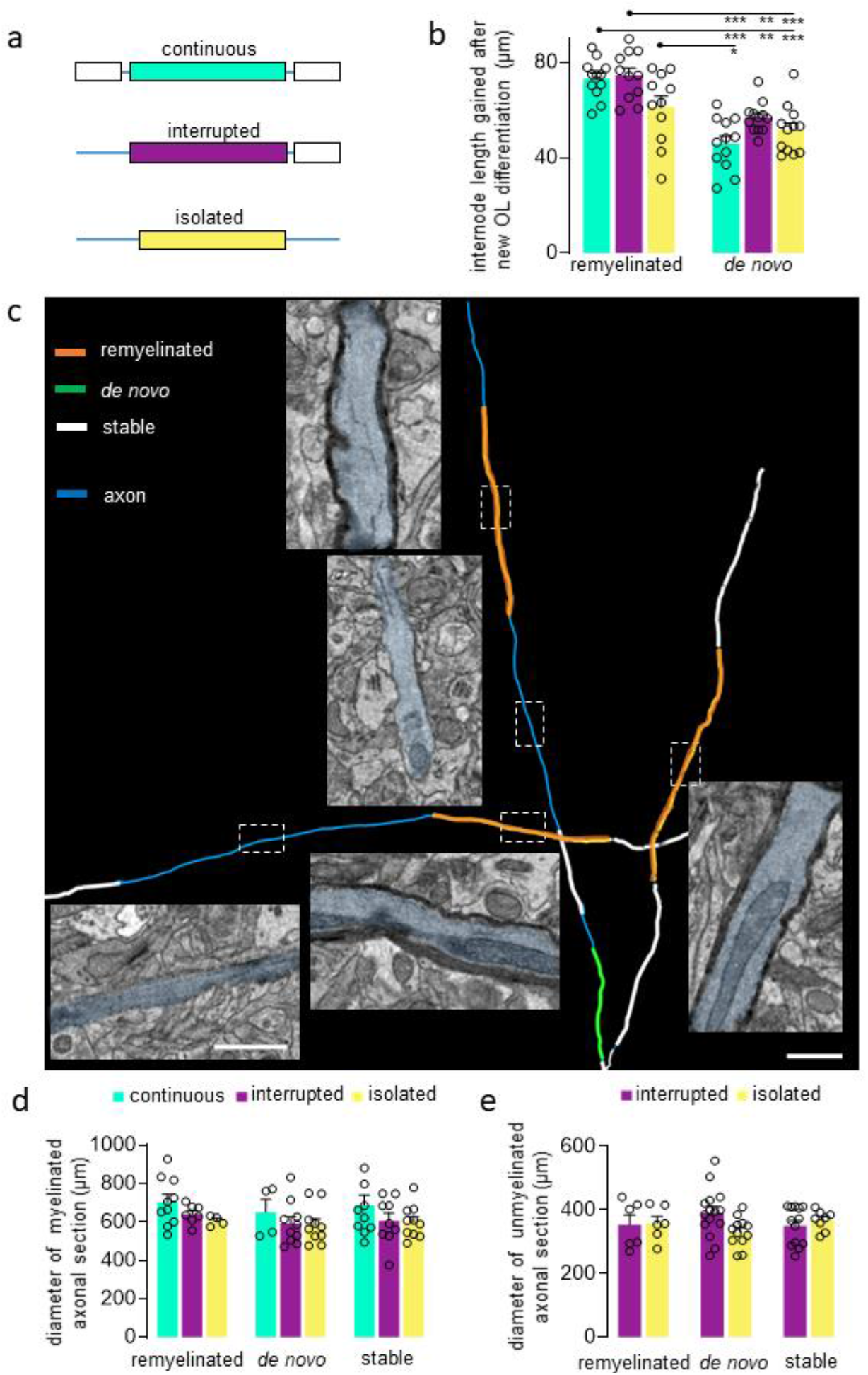
Myelin morphology related to internode type and myelination pattern. **a**: Illustration of the three different internode types in the cortex with the continuous type having two neighboring sheaths, interrupted with an internode on only one side, and isolated without neighboring internodes. **b**: Average internode length of the remyelinated and *de novo* fraction separated within the three internode types (n=12 areas from n=5 animals; mean ± SEM, one-way ANOVA with Tukey’s multiple comparison test). **c**: 3D EM reconstruction of a remyelinated segment from the each internode type displaying similar axonal diameter, myelin thickness and unmyelinated axonal diameter (assessable only for interrupted and isolated segments). **d, e**: Quantification of the axonal diameter of remyelinated *de novo* and stable fraction depending on the internode type in myelinated (**d**: remyelinated n=21, *de novo* n=24, stable n=29; mean ± SEM, one-way ANOVA) and unmyelinated section (**e**: consecutive to remyelinated, n=12, to *de novo* n=27, to stable n=21; mean ± SEM, Kruskal-Wallis with Dunn’s multiple comparisons test). *** P<0.001, **P < 0.01, *P<0.05. Scale bar, electron micrographs: 1μm for all, 3D rendering model, 20 μm.

**Supplementary Figure 7.**
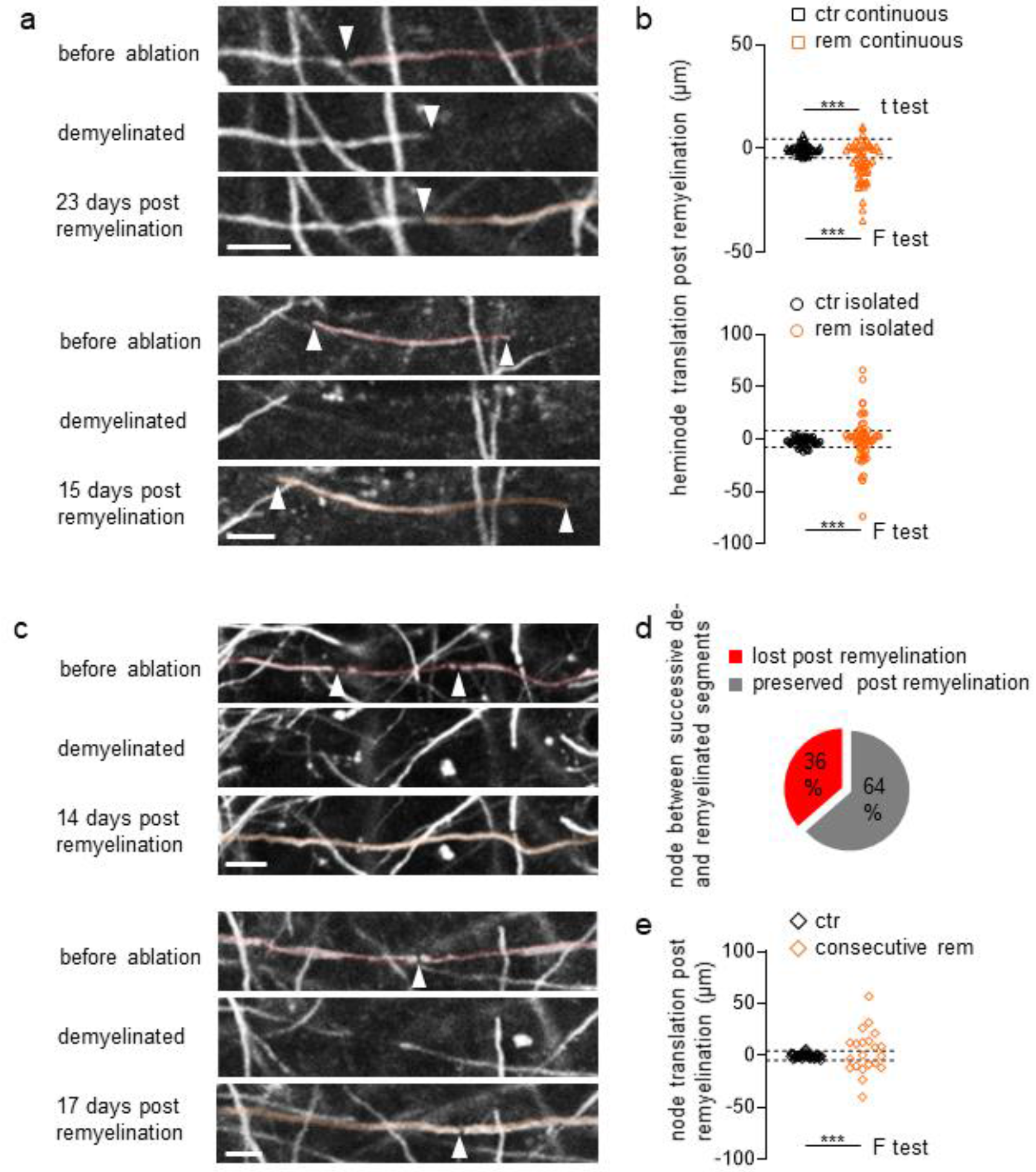
Node positon before and after demyelination-remyelination cycle. **a**: Node repositioning after de- and remyelination for continuous (upper panel) and isolated (lower panel) and isolated myelin sheaths. **b**: Quantification of the heminode misplacement for a remyelinated myelin sheath (continuous: n=46 segments in 5 areas from 5 animals for control and 52/7/5 for remyelinated). Dotted lines plotted at 2 SD of control, i.e. stable nodes (ctr). **c**: Node lost (upper panel) and repositioning (lower panel) between successive de- and remyelinated myelin segments. **d**: Quantification of the lost and repositioned nodes between successive de- and remyelinated myelin segments. Dotted lines plotted at 2xSD of ctr. **e**: Quantification of the node misplacement for successive de- and remyelinated myelin sheath (continuous: n=46 segments in 5 areas from 5 animals for control and 21/10/6 for remyelinated). Statistics: mean ± SEM, t-test and F-test, *** P<0.001. Scale bar: 10μm

**Supplementary Movie 1: Intravital imaging of cortical OL in adult mouse**

Representative area of the somatosensory cortex (layer 1), prior to single OL cell ablation in a *Plp*:GFP mouse, where internode morphology, OL processes and nodes of Ranvier are clearly identifiable. The imaged volume spans 60 μm in z and 203 μm in x/y (2.4 10^6^ μm^3^) and thus exceeds the typical cylindrical volume of a cortical oligodendrocyte in our sample (~ 0.7 10^6^ μm^3^)

## METHODS

### Transgenic animals and virus labeling

Experiments were performed on *Plp*:GFP animals with a C57BL/6 background ^34^, and *Cx_3_cr1*^GFP/+^ knock-in mice ^36^, as well as *Thy1*:GFP^M^ transgenic mice ^39^ bred in our animal facilities. Both female and male animals from 3 to 6 months of age were included in experiments. For viral labeling of OLs, a recombinant, replication deficient AAV (AAV-*Mbp*:mem-tdTom; 10^12^ titer, 0.5μl in Ringer’s solution) was stereotactically injected (coordinates: 2/-2 mm from bregma, 0.2 mm depth) using an ultrathin pulled glass pipette via a 4 mm (diameter) craniotomy performed 14-21 days before the first imaging session.

All animal experiments were performed in accordance with the regulations of the relevant animal welfare acts and protocols approved by the respective regional authorities (Regierung von Oberbayern).

### Cranial window surgery

To gain optical access to cortex for monitoring myelin morphology over time, we performed a craniotomy and implanted a cranial window above the somatosensory cortex of the animals as previously described ^35^. In brief, mice were anaesthetized with a mixture of medetomidin (0.5 mg/kg), midazolam (5 mg/kg) and fentanyl (0.05 mg/kg) intraperitoneally. The craniotomy was performed using a 0.5 mm stainless steel drill head (Meisinger) and a cranial window of 4 mm of diameter was positioned and fixed with dental cement. Mice were given buprenorphine (0.05 - 0.1 mg/kg) for analgesia every 12 h on the days following the surgery.

### *In vivo* imaging

Chronic *in vivo* imaging of the layer 1 OLs and myelin segments was initiated 14-21 days after the implantation of the cranial window and was performed using an Olympus FV-RS microscope (Olympus, Japan) equipped with a femto-second pulsed Ti:Sapphire laser (Mai Tai HP-DS, Spectra-Physics) at a maximum power of 30 mW (measured the back focal plane). Imaging was performed with a resonant scanner using 16x averaging. On the imaging days, animals were anaesthetized starting with 2% isoflurane, placed on the imaging stage and provided with a constant flow of 1.2-1.5% isoflurane for the rest of the imaging period. The physiological state of the animals was continuously monitored with a MouseOx system (Starr Life Science Corp) equipped with a thigh sensor to assess the depth of the anesthesia, oxygenation of the blood and heart rate. On the first imaging day several areas (5-8; x/y/z: 203/203/60 μm^3^) randomly chosen throughout the somatosensory cortex were mapped and imaged with a pixel size of 0.4 μm in x/y and 0.5 μm in z. One OL per non-control area was then ablated by targeting a couple of pixels at the center of the cell (920 nm, 80-120 mW for 1-2 seconds in tornado mode). The area was controlled 2-3 days post ablation to confirm OL death. The same areas were then imaged every 3-7 days for up to 100 days. Control areas were imaged without ablation for the same period. We observed no signs of photo-damage in both ablated and control areas over the imaging period using these imaging conditions. Areas showing deterioration of the imaging quality throughout the chronic imaging were excluded from the analysis.

### Correlative light-to-electron microscopy

After the final imaging session, animals were killed by isoflurane overdose and directly perfused with 4% paraformaldehyde (EMS) and 2.5% glutaraldehyde (EMS) in phosphate buffered saline (EMS). After fixation, the brains where re-positioned under the microscope using the same orientation as set during the *in vivo* imaging. Overview images allowed precise identification of the previously chronically imaged areas. For correlated light and electron microscopy (CLEM), large asymmetric near-infrared branding (NIRB) marks ^63^ were performed 100-200 μm away from the outer most edges of the imaged areas leading to the marking of a large field (x/y: ~1.5*1.5 mm^2^) containing the imaged areas. The tissue was then dissected following these large asymmetric NIRB traces allowing the unambiguous orientation of the areas within the tissue block and further processed for EM imaging.

The *en bloc* staining protocol was adapted from Hua ^64^et al. Briefly, the tissue was initially post-fixed using 2% osmium tetroxide (EMS) in 0.1 M sodium cacodylate (Science Services) buffer (pH 7.4) that was replaced by 2% potassium ferricyanide (Sigma) in the same buffer. After three washing steps in buffer and water the staining was enhanced by reaction with 1% thiocarbohydrazide (Sigma) for 45 min at 50°C. The tissue was washed in water and incubated in 2% aqueous osmium tetroxide (EMS). All osmium incubation steps were carried out over 90 min with substitution by fresh reagents after 45 min, respectively. To intensify the staining further, 2% aqueous uranyl acetate (EMS) was applied overnight at 4°C and subsequently warmed to 50°C for 2 h. The samples were dehydrated in an ascending ethanol series and infiltrated with LX112 (LADD). In order to facilitate trimming and sectioning at an angle parallel to the live imaging plane, the sample was oriented with the pial side parallel to the block tip and cured for 48h.

The block was trimmed by 200 μm at a 90° angle on each side using a TRIM90 diamond knife (Diatome) on an ATUMtome (Powertome, RMC). Consecutive sections were taken with a 35° ultra diamond knife (Diatome) at a nominal cutting thickness of 75 nm and collected on freshly plasma-treated (custom-built, based on Pelco easiGlow, adopted from Mark Terasaki, U. Connecticut), carbon-coated (provided by Richard Schalek and Jeff Lichtman, Harvard U.) Kapton tape ^65^. The area of interest was pre-estimated during sectioning according to endogenous (e.g. blood vessel geometry) and NIRB marks. The corresponding tape region with sections was assembled on the adhesive carbon tape (Science Services) mounted onto 4-inch silicon wafers (Siegert Wafer). Kapton and silicon were connected by adhesive carbon tape strips (Science Services) for grounding.

EM micrographs were acquired on a Crossbeam Gemini 340 SEM (Zeiss) with a four-quadrant backscatter detector at 8 kV. In ATLAS5 Array Tomography (Fibics), the whole wafer area was scanned at 2000-6000 nm/pixel to generate an overview map. In total, 391 sections were selected and imaged at 200*200 nm^2^ and a smaller ROI selected for another imaging round at 50*50 nm^2^. The region of interest was relocated based on this image set compared to the live imaging data. The final volume (370*350*47 μm^3^) was acquired with 6.4 μs dwell time at a voxel size of 12*12*150 nm^3^ (every second section). The high-resolution volumes were generated from several image sets of 35-70*35-85*3 μm^3^, at 3*3*~75 nm^3^ voxel size.

### Image processing and analysis

Post-processing of the presented two-photon imaging was done using the open-source image analysis software, ImageJ/Fiji (http://fiji.sc) ^66^: brightness and contrast were adjusted, maximum projections were performed for display purpose (see figure legend for details), and the time-points represented in the figures were aligned using the StackReg plugin ^67^.

To analyze the position of newly matured OL in ablated conditions the reference point was set at the center of the ablated cell and for the control areas at the center of the imaged area at T0.

To analyze the length and type of myelin sheaths over the ablation experiments, 3D reconstructions of lost and gained myelin segment were performed within the acquired chronic *in vivo* imaging volumes using the simple neurite tracing plugin of Fiji. Internodes ending outside of the imaging area were used in the remyelination efficiency quantification, but excluded from the segment type quantification and length measurements.

The volume covered by an OL in layer 1 of the somatosensory cortex was approximated using the volume of a cylinder of 150μm of diameter (span of the OL in x/y) and 40μm in height (span of the OL in z) in line with previously published data ^19^.

For the 3D EM reconstructions, the images were aligned by a sequence of automatic and manual processing steps in Fiji TrakEM2 ^68^. IMOD software was then used for the manual segmentation, tracing and reconstructions of the CLEM volumes ^69^. G-ratio measurements were performed in three different places along the central section of the internodes. The myelinated axonal diameter measurements were performed in three to five places (depending on the length of the internode) along the entire length of the internode. The measurements of the unmyelinated axonal diameter section was performed in three places within 30 μm of the neighboring myelin heminode. The diameter measurements of the fully unmyelinated axons from the high-resolution EM volumes were performed in three places along the reconstructed axon.

### Statistical analysis

Sample sizes were chosen according to previous *in vivo* imaging studies ^5, 70^. Statistical significance was calculated with Prism (Versions 8.0) using t-test or ANOVA and Tukey’s multiple comparison test (where normal distribution could be assumed by D’Agostino & Pearson or Shapiro-Wilk tests) or Mann-Whitney U-test or Kruskal-Wallis and Dunn’s multiple comparisons test (where non-normal distribution was suspected and confirmed by Shapiro-Wilk test) as described in the figure legends. The cumulative distributions were tested with Kolmogorov-Smirnov test.

### DATA AVAILABILITY

The *Plp*:GFP mouse line was obtained from B. Zalc (ICM, Paris) to whom requests for the mouse line have to be directed. *Cx_3_cr1*^GFP/+^ and *Thy1*-GFP^M^ mice are available from Jackson Laboratory (JAX 005582 and 007788, respectively). The plasmids to generate AAV-*Mbp*:mem-tdTom can be requested from the authors. The primary data from time-lapse imaging and ultrastructural analysis can be requested from the authors.

## REFERENCES

1. Waxman, S. Determinants of conduction velocity in myelinated nerve fibers. Muscle and nerve 3, 141–150 (1980).

2. Simons, M. & Nave, K.A. Oligodendrocytes: myelination and axonal support. Cold Spring Harb Perspect Biol 8, a020479 (2015).

3. Nave, K.A. & Trapp, B.D. Axon-glial signaling and the glial support of axon function. Annu Rev Neurosci 31, 535–561 (2008).

4. O’Brien, J. Stability of the myelin membrane. Science 147, 1099–1107 (1965).

5. Hughes, E.G., Orthmann-Murphy, J.L., Langseth, A.J. & Bergles, D.E. Myelin remodeling through experience-dependent oligodendrogenesis in the adult somatosensory cortex. Nat Neurosci 21, 696–706 (2018).

6. Tomassy, G.S., et al. Distinct profiles of myelin distribution along single axons of pyramidal neurons in the neocortex. Science 344, 319–324 (2014).

7. Bengtsson, S.L., et al. Extensive piano practicing has regionally specific effects on white matter development. Nat Neurosci 8, 1148–1150 (2005).

8. Scholz, J., Klein, M.C., Behrens, T.E. & Johansen-Berg, H. Training induces changes in white-matter architecture. Nat Neurosci 12, 1370–1371 (2009).

9. Zatorre, R.J., Fields, R.D. & Johansen-Berg, H. Plasticity in gray and white: neuroimaging changes in brain structure during learning. Nat Neurosci 15, 528–536 (2012).

10. Sampaio-Baptista, C., et al. Motor skill learning induces changes in white matter microstructure and myelination. J Neurosci 33, 19499–19503 (2013).

11. McKenzie, I., Ohayon, D., Li, H., Paes de Faria, J., Emery, B., Tohyama, K., Richardson, W. D. Motor skill learning requires active central myelination. Science 346 (2014).

12. Micheva, K.D., et al. A large fraction of neocortical myelin ensheathes axons of local inhibitory neurons. Elife 5 (2016).

13. Stedehouder, J., et al. Fast-spiking parvalbumin interneurons are frequently myelinated in the cerebral cortex of mice and humans. Cereb Cortex 27, 5001–5013 (2017).

14. Hill, R.A., Li, A.M. & Grutzendler, J. Lifelong cortical myelin plasticity and age-related degeneration in the live mammalian brain. Nat Neurosci 21, 683–695 (2018).

15. Richardson, W.D., Young, K.M., Tripathi, R.B. & McKenzie, I. NG2-glia as multipotent neural stem cells: fact or fantasy? Neuron 70, 661–673 (2011).

16. Young, K.M., et al. Oligodendrocyte dynamics in the healthy adult CNS: evidence for myelin remodeling. Neuron 77, 873–885 (2013).

17. Fard, M., Van der Meer, F.., Sánchez, P., Cantuti-Castelvetri, L., Mandad, S., Jäkel, S., Fornasiero, E.., Schmitt, S., Ehrlich, M., Starost, L., Kuhlmann, T., Sergiou, C., Schultz, V., Wrzos, C.., Brück, W., Urlaub, H., Dimou, L., Stadelmann, C., Simons, M. BCAS1 expression defines a population of early myelinating oligodendrocytes in multiple sclerosis lesions. Sci. Transl. Med 9 (2017).

18. Yeung, M.S., et al. Dynamics of oligodendrocyte generation and myelination in the human brain. Cell 159, 766–774 (2014).

19. Tripathi, R.B., et al. Remarkable stability of myelinating oligodendrocytes in mice. Cell Rep 21, 316–323 (2017).

20. Ford, M.C., et al. Tuning of Ranvier node and internode properties in myelinated axons to adjust action potential timing. Nat Commun 6, 8073 (2015).

21. Stange-Marten, A., et al. Input timing for spatial processing is precisely tuned via constant synaptic delays and myelination patterns in the auditory brainstem. Proc Natl Acad Sci U S A 114, E4851–E4858 (2017).

22. Gibson, E.M., et al. Neuronal activity promotes oligodendrogenesis and adaptive myelination in the mammalian brain. Science 344, 1252304 (2014).

23. Gledhill, R.F., Harrison, B. M., McDonald W. I.. Pattern of Remyelination in the CNS. Nature 244 (1973).

24. Prineas, J.W., Connell, F.. Remyelination in multiple sclerosis. Ann Neurol 5, 22–31 (1979).

25. Auer, F., Vagionitis, S. & Czopka, T. Evidence for myelin sheath remodeling in the CNS revealed by in vivo imaging. Curr Biol 28, 549–559 e543 (2018).

26. Franklin, R.J. & Ffrench-Constant, C. Remyelination in the CNS: from biology to therapy. Nat Rev Neurosci 9, 839–855 (2008).

27. Gemberling, M., Bailey, T.J., Hyde, D.R. & Poss, K.D. The zebrafish as a model for complex tissue regeneration. Trends Genet 29, 611–620 (2013).

28. Becker, C.G. & Becker, T. Neuronal regeneration from ependymo-radial glial cells: cook, little pot, cook! Dev Cell 32, 516–527 (2015).

29. Calabrese, M., et al. Cortical lesion load associates with progression of disability in multiple sclerosis. Brain 135, 2952–2961 (2012).

30. Calabrese, M., et al. Exploring the origins of grey matter damage in multiple sclerosis. Nat Rev Neurosci 16, 147–158 (2015).

31. Scalfari, A., et al. The cortical damage, early relapses, and onset of the progressive phase in multiple sclerosis. Neurology 90, e2107–e2118 (2018).

32. Chang, A., et al. Cortical remyelination: a new target for repair therapies in multiple sclerosis. Ann Neurol 72, 918–926 (2012).

33. Albert, M., Antel, J., Bruck, W. & Stadelmann, C. Extensive cortical remyelination in patients with chronic multiple sclerosis. Brain Pathol 17, 129–138 (2007).

34. Green, A.J., et al. Clemastine fumarate as a remyelinating therapy for multiple sclerosis (ReBUILD): a randomised, controlled, double-blind, crossover trial. The Lancet 390, 2481–2489 (2017).

35. Cadavid, D., et al. Safety and efficacy of opicinumab in patients with relapsing multiple sclerosis (SYNERGY): a randomised, placebo-controlled, phase 2 trial. The Lancet Neurology 18, 845–856 (2019).

36. Brill, M.S., Lichtman, J.W., Thompson, W., Zuo, Y. & Misgeld, T. Spatial constraints dictate glial territories at murine neuromuscular junctions. J Cell Biol 195, 293–305 (2011).

37. Spassky, N., Goujet-Zalc, C., Parmantier, E., Olivier, C., Martinez, S., Ivanova, A., Ikenaka, K., Macklin, W., Cerruti, I., Zalc, B., Thomas, J. Multiple restricted origin of oligodendrocytes. J. Neurosci 18, 8331–8343 (1998).

38. Holtmaat, A., et al. Long-term, high-resolution imaging in the mouse neocortex through a chronic cranial window. Nat Protoc 4, 1128–1144 (2009).

39. Jung, S., Kreutzberg, G., Aliberti, J., Graemmel, P., Sunshine, M., Sher, A., Littman, D. Analysis of fractalkine receptor CX3CR1 function by targeted deletion and green fluorescent protein reporter gene insertion. Mol and Cell Bio, 4106–4114 (2000).

40. Nimmerjahn A., K.F., Helmchen F. Resting microglial cells are highly dynamic surveillants of brain parenchyma in vivo. Science 308 1314–1318 (2005).

41. Davalos, D., et al. ATP mediates rapid microglial response to local brain injury in vivo. Nat Neurosci 8, 752–758 (2005).

42. Feng, G., Mellor, R., Bernstein, M., Keller-Peck, C., Nguyen, Q., Wallace, M., Nerbonne, J., Lichtman, J., Sanes, J. Imaging Neuronal Subsets in Transgenic Mice Expressing Multiple Spectral Variants of GFP. Neuron 28 (2000).

43. Locatelli, G., et al. Primary oligodendrocyte death does not elicit anti-CNS immunity. Nat Neurosci 15, 543–550 (2012).

44. Drawitsch, F., Karimi, A., Boergens, K., Helmstaedter, M. FluoEM, Virtual labeling of axons in 3-dimensional electron microscopy data for long-range connectomics. eLIFE (2018).

45. Waxman, S., Bennett, M. Relative conduction velocities of small myelinated and non-myelinated fibres in the central nervous system. Nature New Biology 217–219 (1972).

46. Windebank AJ, W.P., Bunge RP, Dyck PJ. Myelination determines the caliber of dorsal root ganglion neurons in culture. J Neurosci., 1563–1569. (1985).

47. Remahl, S., Hildebrand, C. Relations between axons and oligodendroglial cells during initial myelination. I. The glial unit. Journal of Neurocytology 19, 313–328 (1990).

48. Zonouzi, M., et al. Individual oligodendrocytes show bias for inhibitory axons in the neocortex. Cell Rep 27, 2799–2808 e2793 (2019).

49. Makinodan, M., Rosen, K.M., Ito, S. & Corfas, G. A critical period for social experience-dependent oligodendrocyte maturation and myelination. Science 337, 1357–1360 (2012).

50. Xiao, L., et al. Rapid production of new oligodendrocytes is required in the earliest stages of motor-skill learning. Nat Neurosci 19, 1210–1217 (2016).

51. Mitew, S., et al. Pharmacogenetic stimulation of neuronal activity increases myelination in an axon-specific manner. Nat Commun 9, 306 (2018).

52. BØ, L., Vedeler, C., Nyland, H., Trapp, B., MØrk, S. Subpial demyelination in the cerebral cortex of multiple sclerosis patients. Journal of Neuropathology and Experimental Neurology 62 (2003).

53. Miron, V.E., et al. M2 microglia and macrophages drive oligodendrocyte differentiation during CNS remyelination. Nat Neurosci 16, 1211–1218 (2013).

54. Manrique-Hoyos, N., et al. Late motor decline after accomplished remyelination: impact for progressive multiple sclerosis. Ann Neurol 71, 227–244 (2012).

55. Fornasiero, E.F., et al. Precisely measured protein lifetimes in the mouse brain reveal differences across tissues and subcellular fractions. Nat Commun 9, 4230 (2018).

56. Tricaud, N. & Park, H.T. Wallerian demyelination: chronicle of a cellular cataclysm. Cell Mol Life Sci 74, 4049–4057 (2017).

57. Chen, Z. & Trapp, B.D. Microglia and neuroprotection. J Neurochem 136 Suppl 1, 10–17 (2016).

58. Lee, S., et al. A culture system to study oligodendrocyte myelination processes using engineered nanofibers. Nat Methods 9, 917–922 (2012).

59. Baraban, M., Koudelka, S. & Lyons, D.A. Ca (2+) activity signatures of myelin sheath formation and growth in vivo. Nat Neurosci 21, 19–23 (2018).

60. Bechler, M.E., Byrne, L. & Ffrench-Constant, C. CNS Myelin sheath lengths are an intrinsic property of oligodendrocytes. Curr Biol 25, 2411–2416 (2015).

61. Bechler, M.E., Swire, M. & Ffrench-Constant, C. Intrinsic and adaptive myelination-A sequential mechanism for smart wiring in the brain. Dev Neurobiol 78, 68–79 (2018).

## REFERENCES FOR METHODS

34. Spassky, N., Goujet-Zalc, C., Parmantier, E., Olivier, C., Martinez, S., Ivanova, A., Ikenaka, K., Macklin, W., Cerruti, I., Zalc, B., Thomas, J. Multiple restricted origin of oligodendrocytes. J. Neurosci 18, 8331–8343 (1998).

35. Holtmaat, A., et al. Long-term, high-resolution imaging in the mouse neocortex through a chronic cranial window. Nat Protoc 4, 1128–1144 (2009).

36. Jung, S., Kreutzberg, G., Aliberti, J., Graemmel, P., Sunshine, M., Sher, A., Littman, D. Analysis of fractalkine receptor CX3CR1 function by targeted deletion and green fluorescent protein reporter gene insertion. Mol and Cell Bio, 4106–4114 (2000

39. Feng, G., Mellor, R., Bernstein, M., Keller-Peck, C., Nguyen, Q., Wallace, M., Nerbonne, J., Lichtman, J., Sanes, J. Imaging Neuronal Subsets in Transgenic Mice Expressing Multiple Spectral Variants of GFP. Neuron 28 (2000).

63. Bishop, D., et al. Near-infrared branding efficiently correlates light and electron microscopy. Nat Methods 8, 568–570 (2011).

64. Hua, Y., Laserstein, P. & Helmstaedter, M. Large-volume en-bloc staining for electron microscopy-based connectomics. Nat Commun 6, 7923 (2015).

65. Kasthuri, N., et al. Saturated Reconstruction of a Volume of Neocortex. Cell 162, 648–661 (2015).

66. Schindelin, J., et al. Fiji: an open-source platform for biological-image analysis. Nat Methods 9, 676–682 (2012).

67. Thévenaz P, R.U., Unser M. A pyramid approach to subpixel registration based on intensity. IEEE Trans Image Process 7, 27–24.

68. Cardona, A., et al. TrakEM2 software for neural circuit reconstruction. PLoS One 7, e38011 (2012).

69. Kremer JR, M.D., McIntosh JR. Computer visualization of three-dimensional image data using IMOD. J Struct Biol. 116, 71–76. (1996).

70. Nikic, I., et al. A reversible form of axon damage in experimental autoimmune encephalomyelitis and multiple sclerosis. Nat Med 17, 495–499 (2011).

